# An alternative route for β-hydroxybutyrate metabolism supports fatty acid synthesis in cancer cells

**DOI:** 10.1101/2024.10.31.621317

**Authors:** Faith C. Kaluba, Thomas J. Rogers, Yu-Jin Jeong, Althea Waldhart, Kelly H. Sokol, Cameron J. Lee, Samuel R. Daniels, Joseph Longo, Amy Johnson, Ryan D. Sheldon, Russell G. Jones, Evan C. Lien

**Affiliations:** Department of Metabolism and Nutritional Programming, Van Andel Institute, 333 Bostwick Ave. NE, Grand Rapids, MI 49503; Mass Spectrometry Core, Van Andel Institute, 333 Bostwick Ave. NE, Grand Rapids, MI 49503; Van Andel Institute Graduate School, 333 Bostwick Ave. NE, Grand Rapids, MI 49503

**Author notes:** These authors contributed equally to this work (Faith C. Kaluba, Thomas J. Rogers).

## Abstract

Cancer cells are exposed to diverse metabolites in the tumor microenvironment that are used to support the synthesis of nucleotides, amino acids, and lipids needed for rapid cell proliferation^1–3^. Recent work has shown that ketone bodies such as β-hydroxybutyrate (β-OHB), which are elevated in circulation under fasting conditions or low glycemic diets, can serve as an alternative fuel that is metabolized in the mitochondria to provide acetyl-CoA for the tricarboxylic acid (TCA) cycle in some tumors^4–7^. Here, we discover a non-canonical route for β-OHB metabolism, in which β-OHB can bypass the TCA cycle to generate cytosolic acetyl-CoA for *de novo* fatty acid synthesis in cancer cells. We show that β-OHB-derived acetoacetate in the mitochondria can be shunted into the cytosol, where acetoacetyl-CoA synthetase (AACS) and thiolase convert it into acetyl-CoA for fatty acid synthesis. This alternative metabolic routing of β-OHB allows it to avoid oxidation in the mitochondria and net contribute to anabolic biosynthetic processes. In cancer cells, β-OHB is used for fatty acid synthesis to support cell proliferation under lipid-limited conditions *in vitro* and contributes to tumor growth under lipid-limited conditions induced by a calorie-restricted diet *in vivo*. Together, these data demonstrate that β-OHB is preferentially used for fatty acid synthesis in cancer cells to support tumor growth.

## Main Text

Under fasting conditions, insulin resistance, or low glycemic diets (such as the ketogenic diet or caloric restriction) that decrease blood glucose levels, ketogenesis occurs predominantly in the liver to produce ketone bodies from fatty acids and ketogenic amino acids. Ketone body oxidation then becomes a major contributor to energy metabolism for extrahepatic tissues, thereby sparing glucose^8^. We and others recently demonstrated that certain cancer cell lines can also metabolize β-OHB through the TCA cycle^4,5,7^. β-OHB oxidation occurs in the mitochondria, where β-OHB dehydrogenase 1 (BDH1) and 3-oxoacid CoA-transferase 1 (OXCT1) are localized. BDH1, OXCT1, and mitochondrial thiolase convert β-OHB into acetyl-CoA, which subsequently enters the TCA cycle to contribute to energy production through oxidative phosphorylation (Fig. 1A). We assessed protein levels of BDH1 and OXCT1 in a panel of cancer cell lines and found three cell lines that express both enzymes: AL1376, a pancreatic ductal adenocarcinoma (PDAC) cell line derived from the LSL-*Kras*(G12D/+);*Trp53*(fl/fl);Pdx1-Cre mouse PDAC model^4,9^; B16, a mouse-derived melanoma cell line; and MIA PaCa-2, a human PDAC cell line (Fig. 1B). Even though β-OHB is considered to be an alternative to glucose for fueling energy production in tissues, we found that β-OHB minimally rescued the proliferation defect of glucose-starved cells (Fig. 1C), suggesting that β-OHB does not replace glucose to support cell proliferation. β-OHB also contributes to *de novo* fatty acid synthesis through the production of mitochondrial citrate, which is subsequently exported into the cytosol to generate the cytosolic acetyl-CoA needed for fatty acid synthesis (Fig. 1A). Because fatty acid synthesis is required for cancer cells to proliferate when extracellular lipid levels are limiting^10^, we asked whether β-OHB rescues the proliferation defect of cells cultured in lipid-depleted media. Indeed, β-OHB promoted cell proliferation in lipid-depleted media across all three cell lines (Fig. 1D), and treating cells with the fatty acid synthase (FASN) inhibitor GSK2194069 prevented this rescue (Fig. 1E). Taken together, these data indicate that β-OHB can promote the proliferation of cancer cells under conditions of extracellular lipid limitation by contributing to *de novo* fatty acid synthesis.

**Figure 1.**
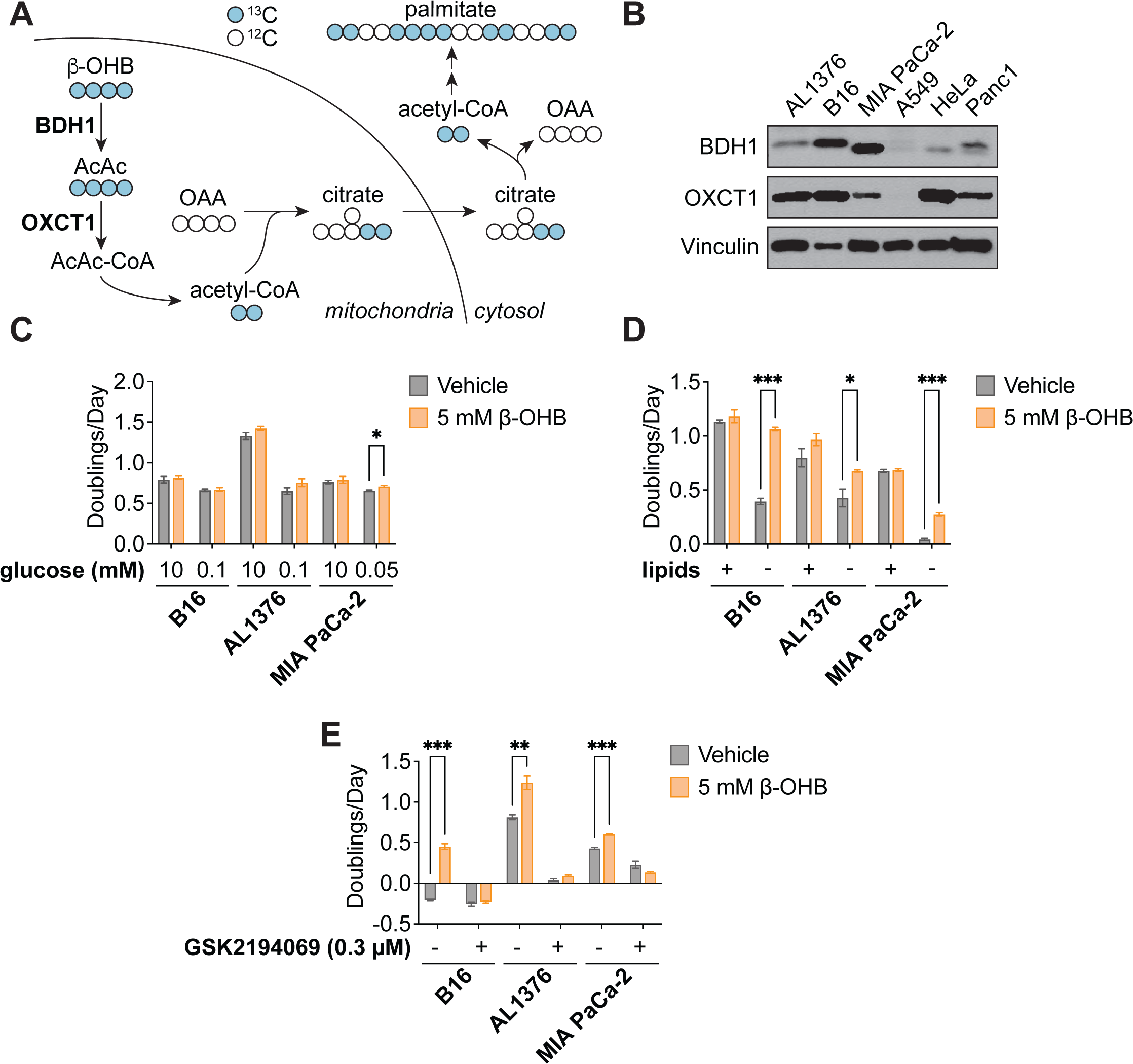
β-OHB promotes the proliferation of lipid-starved cancer cells. **A**, Schematic of β-OHB metabolism and ^13^C labeling derived from [U-^13^C]-β-OHB. AcAc, acetoacetate; BDH1, β-OHB dehydrogenase 1; OXCT1, 3-oxoacid CoA-transferase 1; OAA, oxaloacetate. **B**, Immunoblot for BDH1, OXCT1, and vinculin in the indicated cancer cell lines. **C**, Proliferation rates of the indicated cancer cell lines grown in high or low glucose conditions, with or without 5 mM β-OHB. **D**, Proliferation rates of the indicated cancer cell lines grown in lipid-replete versus lipid-depleted culture media, with or without 5 mM β-OHB. **E**, Proliferation rates of the indicated cancer cell lines grown in lipid-depleted media, with or without 5 mM β-OHB and 0.3 µM of the FASN inhibitor GSK2194069. Data are presented as mean ± s.e.m; *n* = 3 biologically independent replicates. Comparisons were made using a two-tailed Student’s t test (**C-E**). \**P<0.05*, \*\**P<0.01*, \*\*\**P<0.001*.

To confirm that β-OHB is used to synthesize fatty acids such as palmitate, we labeled each cell line with [U-^13^C]-β-OHB for 48 hours and assessed palmitate labeling by gas chromatography-mass spectrometry (GC-MS). In all three cell lines, we detected substantial labeling of palmitate by [U-^13^C]-β-OHB in both lipid-replete and lipid-depleted media (Fig. 2A-C). We also observed substantial ^13^C labeling in citrate, the precursor of cytosolic acetyl-CoA (Fig. 2D-F). By using isotopomer spectral analysis (ISA) through FAMetA^11^, we also calculated the enrichment of β-OHB-derived carbons in cytosolic acetyl-CoA (Fig. 2D-F). We found that cytosolic acetyl-CoA was highly labeled by β-OHB, and in some cases (such as for AL1376 and MIA PaCa-2 cells), it was labeled to a greater extent than citrate (Fig. 2D-F). This was surprising because labeling of downstream metabolites typically becomes more diluted, rather than enriched, compared to the labeling of their upstream precursors. As a comparison, we also labeled each cell line with [U-^13^C]-glucose for 48 hours. Glucose also labeled palmitate, citrate, and cytosolic acetyl-CoA, but in this case cytosolic acetyl-CoA labeling was diluted relative to citrate labeling in all three cell lines (Fig. 2G-H, Extended Data Fig. 1A-D). These results indicate that higher labeling of cytosolic acetyl-CoA relative to citrate is unique to β-OHB and suggests that β-OHB may be preferentially used over glucose for fatty acid synthesis. Indeed, we found that the presence of unlabeled β-OHB strongly suppressed [U-^13^C]-glucose labeling of palmitate, citrate, and cytosolic acetyl-CoA in all three cell lines (Fig. 2I-J, Extended Data Fig. 1E-H). To quantify the amount of ^13^C label dilution versus enrichment in cytosolic acetyl-CoA relative to citrate, we calculated the fold change in labeled cytosolic acetyl-CoA compared to labeled citrate (Fig. 2K). This confirmed that [U-^13^C]-β-OHB highly labeled cytosolic acetyl-CoA, in some cases more than it labeled citrate, whereas [U-^13^C]-glucose labeling of cytosolic acetyl-CoA was always diluted compared to citrate. Moreover, the presence of unlabeled β-OHB further diluted cytosolic acetyl-CoA labeling by [U-^13^C]-glucose. Finally, we noted that cytosolic acetyl-CoA is also used for fatty acid elongation, which includes the elongation of polyunsaturated fatty acids (PUFAs) to generate long-chain and very long-chain PUFAs such as 20:3(n-6), 20:4(n-6), 22:4(n-6), and 22:6(n-3). Indeed, we found that both [U-^13^C]-β-OHB and [U-^13^C]-glucose tracing led to [M+2] labeling of these PUFAs in all three cell lines, and importantly, the presence of unlabeled β-OHB suppressed the labeling of these PUFAs by [U-^13^C]-glucose (Extended Data Fig. 2A-C). Together, these data indicate that β-OHB is preferred as a substrate for fatty acid synthesis and elongation compared to glucose.

**Figure 2.**
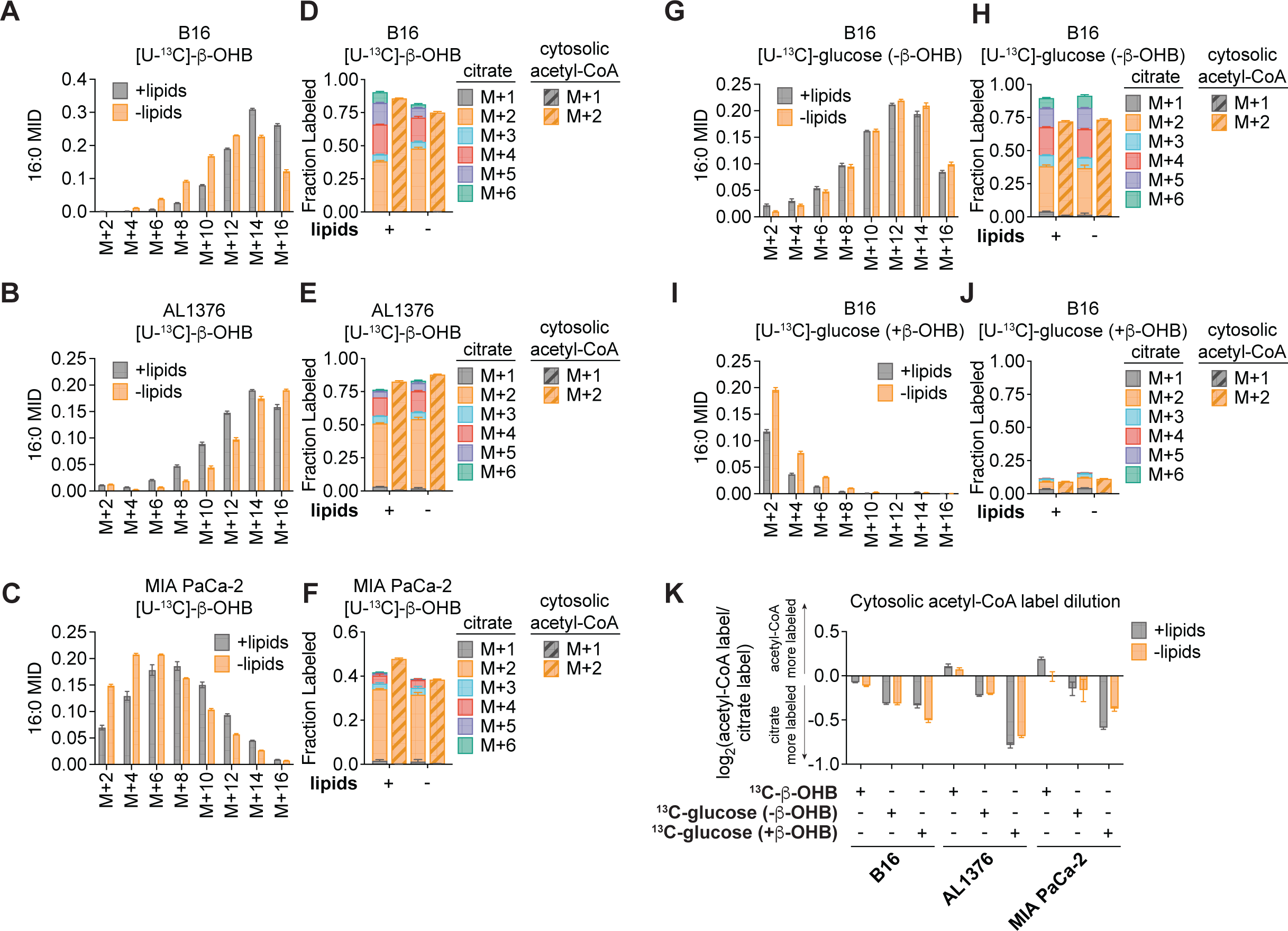
β-OHB is preferentially used over glucose for fatty acid synthesis. **A-C**, 16:0 mass isotopomer distribution (MID) for B16 (**A**), AL1376 (**B**), and MIA PaCa-2 (**C**) cells labeled with [U-^13^C]-β-OHB for 48 h in lipid-replete versus lipid-depleted culture media. **D-F**, Citrate MID (solid bars) and cytosolic acetyl-CoA MID (dashed bars) for B16 (**D**), AL1376 (**E**), and MIA PaCa-2 (**F**) cells labeled with [U-^13^C]-β-OHB for 48 h in lipid-replete versus lipid-depleted culture media. **G-J**, 16:0 MID (**G**, **I**) and citrate and cytosolic acetyl-CoA MID (**H**, **J**) for B16 cells labeled with [U-^13^C]-glucose for 48 h in lipid-replete versus lipid-depleted culture media, either without unlabeled β-OHB (**G**, **H**) or with 5 mM unlabeled β-OHB (**I**, **J**). **K**, Cytosolic acetyl-CoA label dilution from citrate, as calculated by the log_2_(fold change) of the total fraction of cytosolic acetyl-CoA labeled versus the total fraction of citrate labeled, in B16, AL1376, and MIA PaCa-2 cells under the indicated tracing conditions. Data are presented as mean ± s.e.m; *n* = 3 biologically independent replicates.

We hypothesized that higher labeling of cytosolic acetyl-CoA relative to citrate can only occur if β-OHB is converted to cytosolic acetyl-CoA through a citrate-independent route. To test this, we used CRISPR-Cas9 to knock out either *Bdh1* or *Oxct1* in B16 cells, the cell line in which β-OHB rescue of cell proliferation in lipid-depleted conditions was most robust (Fig. 1D). *Bdh1* or *Oxct1* knockout was confirmed by Western blot (Fig. 3A, D). In *Bdh1* knockout cells, β-OHB no longer rescued cell proliferation in lipid-depleted media, suggesting that BDH1 is required for the synthesis of palmitate from β-OHB (Extended Data Fig. 3A-B). Indeed, tracing with [U-^13^C]-β-OHB revealed markedly reduced labeling of citrate and cytosolic acetyl-CoA in *Bdh1* knockout cells (Fig. 3B). This was also reflected by a shift to the left in the mass isotopomer distribution (MID) of labeled palmitate (Fig. 3C). *Bdh1* loss also suppressed [M+2] labeling of the omega-6 PUFAs 20:3(n-6), 20:4(n-6), and 22:4(n-6) (Extended Data Fig. 4A). Residual labeling of citrate, cytosolic acetyl-CoA, palmitate, and PUFAs in *Bdh1* knockout cells suggests that there may be additional enzyme(s) with β-OHB dehydrogenase activity. In contrast, β-OHB still partially rescued the proliferation of *Oxct1* knockout cells in lipid-depleted media (Extended Data Fig. 3C), indicating that OXCT1 is not the only route for the synthesis of palmitate from β-OHB. In these cells, labeling of citrate from [U-^13^C]-β-OHB was diminished to the same extent as in *Bdh1* knockout cells, as expected (Fig. 3E). However, β-OHB still substantially labeled cytosolic acetyl-CoA (Fig. 3E), reflected by a less robust shift to the left in the MID of palmitate in the *Oxct1* knockout cells compared to the *Bdh1* knockout cells (compare Fig. 3F to Fig. 3C). Consistently, *Oxct1* loss partially suppressed [M+2] PUFA labeling, but not to the same extent as *Bdh1* loss (Extended Data Fig. 4A). These results confirm that β-OHB can bypass OXCT1-dependent mitochondrial citrate production to generate the cytosolic acetyl-CoA needed for fatty acid synthesis and elongation.

**Figure 3.**
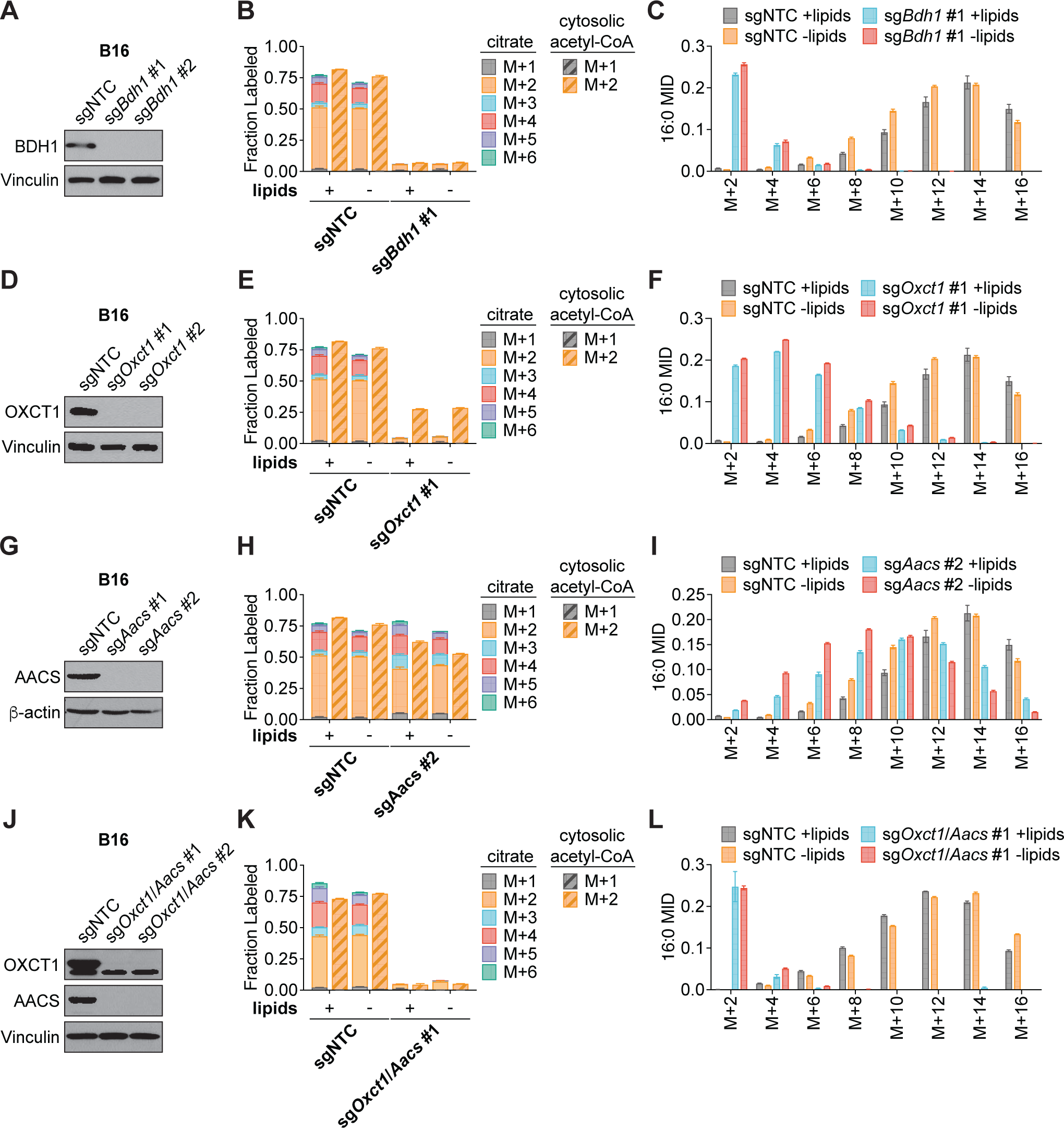
β-OHB can contribute to fatty acid synthesis through a citrate-independent route that requires AACS. **A**, Immunoblot for BDH1 and vinculin in the indicated B16 knockout lines. NTC, non-targeting control. **B**, **C**, Citrate (solid bars) and cytosolic acetyl-CoA (dashed bars) MIDs (**B**), and 16:0 MID (**C**) in B16 sgNTC versus sg*Bdh1* #1 cells labeled with [U-^13^C]-β-OHB for 48 h in lipid-replete versus lipid-depleted culture media. **D**, Immunoblot for OXCT1 and vinculin in the indicated B16 knockout lines. **E**, **F**, Citrate (solid bars) and cytosolic acetyl-CoA (dashed bars) MIDs (**E**), and 16:0 MID (**F**) in B16 sgNTC versus sg*Oxct1* #1 cells labeled with [U-^13^C]-β-OHB for 48 h in lipid-replete versus lipid-depleted culture media. **G**, Immunoblot for AACS and β-actin in the indicated B16 knockout lines. **H**, **I**, Citrate (solid bars) and cytosolic acetyl-CoA (dashed bars) MIDs (**H**), and 16:0 MID (**I**) in B16 sgNTC versus sg*Aacs* #2 cells labeled with [U-^13^C]-β-OHB for 48 h in lipid-replete versus lipid-depleted culture media. **J**, Immunoblot for OXCT1, AACS, and vinculin in the indicated B16 knockout lines. **K**, **L**, Citrate (solid bars) and cytosolic acetyl-CoA (dashed bars) MIDs (**K**), and 16:0 MID (**L**) in B16 sgNTC versus sg*Oxct1*/*Aacs* #1 cells labeled with [U-^13^C]-β-OHB for 48 h in lipid-replete versus lipid-depleted culture media. Data are presented as mean ± s.e.m; *n* = 3 biologically independent replicates.

Based on these results, we reasoned that mitochondrial acetoacetate (the substrate for OXCT1) is likely transported into the cytosol to be used for cytosolic acetyl-CoA production. Acetoacetyl-CoA synthetase (AACS) is a cytosolic enzyme that produces acetoacetyl-CoA from acetoacetate, which can then be cleaved by acetoacetyl-CoA thiolase to produce cytosolic acetyl-CoA^12^. AACS is highly expressed in kidney, heart, brain, adipose tissue, and osteoclasts^13–15^, and metabolic tracer studies with ^14^C-labeled β-OHB have provided evidence of cytosolic activation of acetoacetate and its subsequent contribution to fatty acid and cholesterol synthesis in rat livers, brain, spinal cord, skin, and lactating mammary glands^16–19^. We therefore asked whether AACS can also contribute to palmitate synthesis from β-OHB in cancer cells. Using CRISPR-Cas9, we knocked out *Aacs* in B16 cells (Fig. 3G) and found that the ability of β-OHB to rescue the proliferation of these cells in lipid-depleted media was partially blunted in *Aacs* knockout cells (Extended Data Fig. 3D). Labeling of citrate from [U-^13^C]-β-OHB was unaffected by the loss of *Aacs*, confirming that mitochondrial β-OHB metabolism through the TCA cycle remained intact (Fig. 3H). However, *Aacs* loss reduced labeling of cytosolic acetyl-CoA (Fig. 3H) and caused a shift to the left in the MID of palmitate (Fig. 3I). Labeling of cytosolic acetyl-CoA was also diluted compared to citrate labeling in *Aacs* knockout cells (Fig. 3H), consistent with the model that *Aacs* knockout abolishes citrate-independent production of cytosolic acetyl-CoA from β-OHB. Finally, *Aacs* loss also partially impaired [M+2] labeling of PUFAs (Extended Data Fig. 4A). These results indicate that β-OHB is preferentially used for fatty acid synthesis and elongation through two routes: a citrate-dependent route that requires OXCT1, and a citrate-independent route that requires AACS.

Of note, the citrate-dependent route appears to be the more dominant route for β-OHB-derived fatty acid synthesis in B16 cells because the loss of cytosolic acetyl-CoA labeling, the shift to the left in the palmitate MID, and the loss of PUFA labeling was greater in *Oxct1* knockout cells compared to *Aacs* knockout cells (compare Fig. 3H with Fig. 3E, and Fig. 3I with Fig. 3F; Extended Data Fig. 4A). Whether the citrate-independent AACS route might be more dominant in other cancer cell types, and whether the relative activities of these two routes might be regulated in certain contexts, remains to be determined. To confirm that β-OHB utilization for fatty acid synthesis occurs primarily through these two routes, we used CRISPR-Cas9 to knock out both *Oxct1* and *Aacs* in B16 cells (Fig. 3J). Loss of both genes substantially reduced labeling of citrate, cytosolic acetyl-CoA, and PUFAs by [U-^13^C]-β-OHB to the same extent as *Bdh1* loss (Fig. 3K, Extended Data Fig. 4B). This was also reflected by a robust shift to the left in the MID of labeled palmitate comparable to that observed in *Bdh1* knockout cells (Fig. 3L). These results collectively confirm that both OXCT1 and AACS are required for the contribution of β-OHB to fatty acid synthesis and elongation.

We next asked whether β-OHB is preferentially used for fatty acid synthesis in tumors *in vivo*. C57BL/6J mice were subcutaneously implanted with control B16 sgNTC cells on one flank and with either B16 sg*Bdh1* #1, sg*Oxct1* #1, sg*Aacs* #2, or sg*Oxct1*/*Aacs* #1 cells on the other flank. After tumors formed, mice were fasted for 16 hours before being infused with [U-^13^C]-β-OHB through a tail vein catheter with a priming bolus of 478 mg/kg for 1 minute followed by a constant infusion at 9.75 mg/kg/min for 6.5 hours. Using this approach, we were able to achieve ∼50% ^13^C label enrichment in plasma β-OHB (Fig. 4A). Interestingly, while [M+4] β-OHB was the predominant isotopomer in the plasma, a smaller fraction was [M+2] labeled (Fig. 4A), suggesting that ketogenic tissues (e.g., the liver) may be breaking down uniformly labeled [M+4] β-OHB into [M+2] acetyl-CoA, which was then combined with unlabeled acetyl-CoA to generate [M+2] β-OHB that was released into circulation. Importantly, palmitate was minimally labeled in the plasma (Extended Data Fig. 5A), which suggests that any palmitate labeling in tumors likely resulted from tumor β-OHB metabolism.

**Figure 4.**
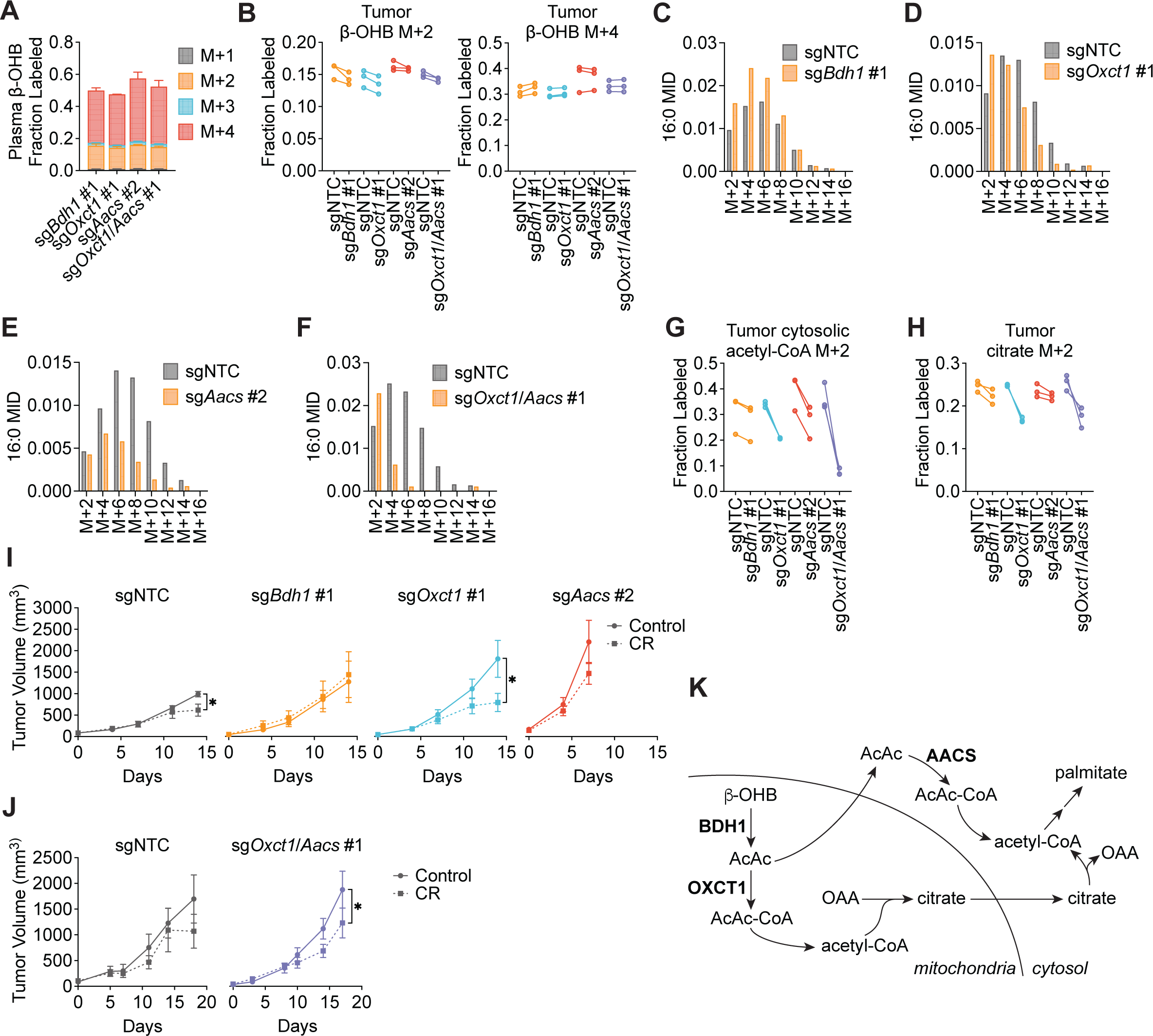
β-OHB contributes to fatty acid synthesis through both OXCT1 and AACS in tumors *in vivo*. C57BL/6J mice bearing a B16 sgNTC tumor on one flank and a knockout tumor on the other flank were infused with [U-^13^C]-β-OHB for 6.5 h. **A**, Plasma β-OHB labeling in mice bearing the indicated knockout tumors. **B**, [M+2] and [M+4] fractional labeling of β-OHB in the indicated sgNTC versus knockout tumors. Data are paired between sgNTC and knockout tumors from the same mouse. n = 3 biological replicates for each knockout tumor. **C-F**, Representative 16:0 MIDs from n = 1 biological replicate of a mouse bearing an sgNTC tumor versus a sg*Bdh1* #1 tumor (**C**), sg*Oxct1* #1 tumor (**D**), sg*Aacs* #2 tumor (**E**), and sg*Oxct1*/*Aacs* #1 tumor (**F**). See Extended Data Fig. 5 for additional biological replicates. **G**, **H**, [M+2] fractional labeling of cytosolic acetyl-CoA (**G**) and citrate (**H**) in the indicated sgNTC versus knockout tumors. Data are paired between sgNTC and knockout tumors from the same mouse. n = 3 biological replicates for each knockout tumor. **I**, **J**, Tumor volumes of subcutaneous B16 sgNTC, sg*Bdh1* #1, sg*Oxct1* #1, sg*Aacs* #2 (**I**), and sg*Oxct1*/*Aacs* #1 (**J**) allografts in male mice exposed to a control or caloric restriction (CR) diet. (**I**) sgNTC Control n = 6 mice, CR n = 6 mice; sg*Bdh1* #1 Control n = 6 mice, CR n = 6 mice; sg*Oxct1* #1 Control n = 5 mice, CR n = 5 mice; sg*Aacs* #2 Control n = 6 mice, CR n = 6 mice. (**J**) sgNTC Control n = 3 mice, CR n = 4 mice; sg*Oxct1*/*Aacs* #1 Control n = 4 mice, CR n = 4 mice. **K**, Schematic of β-OHB contribution to fatty acid synthesis through both a mitochondrial citrate-dependent route via OXCT1 and a citrate-independent route via AACS. Data are presented as mean ± s.e.m. Comparisons were made using a two-way ANOVA (**I, J**). \**P<0.05*.

In tumors, we observed both [M+4] and [M+2] β-OHB labeling for a combined enrichment of ∼50% (Fig. 4B). Within each mouse, [U-^13^C]-β-OHB enrichment was similar between the sgNTC tumor on one flank versus the knockout tumor on the other flank, which enabled direct comparison of labeling patterns in metabolites downstream of β-OHB between the two tumors (Fig. 4B). Surprisingly, in contrast to our *in vitro* results, *Bdh1* loss did not cause a robust shift to the left in the palmitate MID compared to sgNTC tumors (Fig. 4C, Extended Data Fig. 5B). This was reflected by a minimal reduction in [M+2] cytosolic acetyl-CoA and [M+2] citrate labeling in sg*Bdh1* tumors (Fig. 4G, H). One possible explanation for this may be that the [U-^13^C]-β-OHB infusion could also label circulating acetoacetate in the plasma, which can be taken up by tumors and bypass BDH1 to synthesize palmitate through either OXCT1 or AACS. Alternatively, it is possible that another enzyme with β-OHB dehydrogenase activity, such as BDH2, may compensate for BDH1 loss in tumors *in vivo*. In contrast, we observed a stronger shift to the left in the palmitate MID in sg*Oxct1* and sg*Aacs* tumors, and therefore a more robust decrease in cytosolic acetyl-CoA labeling (Fig. 4D-E, 4G, Extended Data Fig. 5C-D). As expected, *Oxct1* loss, but not *Aacs* loss, resulted in decreased tumor [M+2] citrate labeling (Fig. 4H). Finally, loss of both *Oxct1* and *Aacs* in tumors caused the strongest shift to the left in the palmitate MID and an almost complete loss in cytosolic acetyl-CoA labeling, compared to the single knockouts (Fig. 4F-G, Extended Data Fig. 5E). sg*Oxct1*/*Aacs* tumors also had decreased [M+2] citrate labeling (Fig. 4H). Collectively, these results demonstrate that β-OHB utilization for fatty acid synthesis in tumors *in vivo* also occurs through both the mitochondrial citrate-dependent OXCT1 route and the cytosolic citrate-independent AACS route.

To determine whether a similar labeling pattern was observed with β-OHB contribution to PUFA elongation in tumors, we measured [M+2] labeling of 20:3(n-6), 20:4(n-6), and 22:4(n-6). Interestingly, we found distinct effects of each gene knockout on different PUFAs. Labeling of 20:3(n-6) was reduced in all knockout tumors, and this reduction was most dramatic in sg*Aacs* and sg*Oxct1*/*Aacs* tumors (Extended Data Fig. 5F). In contrast, 20:4(n-6) labeling was reduced only in sg*Oxct1*/*Aacs* tumors (Extended Data Fig. 5G), and 22:4(n-6) labeling was elevated in sg*Bdh1* and sg*Oxct1* tumors, but reduced in sg*Aacs* and sg*Oxct1*/*Aacs* tumors (Extended Data Fig. 5H). This differs from the PUFA tracing patterns observed *in vitro*, where loss of *Bdh1*, *Oxct1*, or *Aacs* all reduced PUFA labeling by β-OHB (Extended Data Fig. 4). These results raise the intriguing possibility that in tumors *in vivo*, acetyl-CoA derived from the AACS route may be preferentially used for PUFA elongation. How cells might differentially use acetyl-CoA derived from the OXCT1 route versus the AACS route will require further study.

Finally, we asked how these pathways might influence tumor growth. We previously demonstrated that caloric restriction (CR) decreases lipid levels and increases β-OHB levels in the tumor microenvironment^4^. Therefore, we hypothesized that under these conditions, the contribution of β-OHB to fatty acid synthesis might be important for sustaining tumor growth. To test this, we subcutaneously implanted C57BL6/J mice with B16 sgNTC, sg*Bdh1* #1, sg*Oxct1* #1, or sg*Aacs* #2 cells. After palpable tumors formed, mice were fed with either a control diet or CR. We found that the growth of sgNTC control B16 tumors was only slightly impaired by CR (Fig. 4I). sg*Bdh1* tumors were unaffected by CR (Fig. 4I), consistent with minimal loss of palmitate labeling by [U-^13^C]-β-OHB in sg*Bdh1* tumors in our in vivo labeling experiments (Fig. 4C, Extended Data Fig. 5B). In contrast, the growth of sg*Oxct1* tumors was significantly inhibited by CR (Fig. 4I), which correlates with decreased palmitate labeling by [U-^13^C]-β-OHB in these tumors (Fig. 4D, Extended Data Fig. 5C). The growth of sg*Aacs* tumors in calorically restricted mice trended towards a decrease, but was not statistically significant (Fig. 4I). The stronger effect of CR on inhibiting sg*Oxct1* versus sg*Aacs* tumors might be explained by our finding that the citrate-dependent OXCT1 route is the more dominant route for β-OHB-derived fatty acid synthesis in B16 cells (Fig. 3); however, the interpretation of the data for the sg*Aacs* tumors may also be confounded by how these tumors reached endpoint size in both diet groups by 7 days after diet administration, in contrast to 14 days for the sgNTC, sg*Bdh1*, and sg*Oxct1* tumors. In fact, we were surprised to find that the loss of any of the β-OHB metabolism genes accelerated B16 tumor growth in mice fed a control diet, and this accelerated growth was particularly pronounced for both *Oxct1* and *Aacs* loss (Fig. 4I). This result might suggest that β-OHB metabolism may actually restrain the growth of B16 tumors in mice fed a control diet. Because loss of both *Oxct1* and *Aacs* reduced palmitate and cytosolic acetyl-CoA labeling to a stronger extent than the knockout of either gene alone (Fig. 4F-G, Extended Data Fig. 5E), we also asked whether the growth of sg*Oxct1*/*Aacs* tumors might be more impaired by CR. We found that CR inhibited the growth of sg*Oxct1*/*Aacs* tumors (Fig. 4J), but not to a greater extent than that observed for sg*Oxct1* tumors (Fig. 4I). Taken together, these results suggest that β-OHB utilization for fatty acid synthesis through OXCT1 and AACS has a minor contribution to sustaining B16 tumor growth in the context of a CR diet. Whether this pathway might have a greater contribution to tumor growth under other contexts will require further study.

In summary, in this study we demonstrate that cancer cells capable of metabolizing β-OHB preferentially use it for cytosolic acetyl-CoA production to support the *de novo* synthesis of fatty acids. In addition to the canonical route of cytosolic acetyl-CoA synthesis through OXCT1-dependent production of citrate from the TCA cycle, our data show that β-OHB-derived acetoacetate in mitochondria can be exported to the cytosol, where it is then converted into cytosolic acetyl-CoA through AACS and cytosolic thiolases (Fig. 4K). Notably, in contrast to other fuels such as glucose and glutamine, β-OHB is not an anaplerotic substrate that can net contribute carbons to the TCA cycle, raising the question of how β-OHB can be an effective alternative fuel used by cancer cells for biomass production. Our work shows that by bypassing TCA cycle-dependent mitochondrial citrate production, AACS-dependent production of cytosolic acetyl-CoA allows β-OHB to avoid its oxidation and instead be channeled toward the synthesis of fatty acids needed to support rapid cancer cell proliferation. This feature distinguishes β-OHB from glucose and glutamine as a lipogenic substrate since carbons from glucose and glutamine need to be routed through citrate production to produce cytosolic acetyl-CoA. In this sense, β-OHB is similar to acetate, which has also been proposed as an alternative fuel for tumors that contributes to lipogenesis by being directly converted into cytosolic acetyl-CoA via ACSS2^20,21^. Finally, we note that cytosolic acetyl-CoA has additional fates beyond fatty acid synthesis, including cholesterol synthesis^22^ and histone acetylation^23,24^. Our identification of an alternative pathway for β-OHB metabolism that channels β-OHB toward cytosolic acetyl-CoA production will help to better define how ketone body metabolism influences diverse downstream cellular processes to alter cancer progression.

## Acknowledgements

We thank members of the Lien laboratory and Russell Jones’ laboratory for useful discussions. We also thank Lisa DeCamp for assistance with *in vivo* stable isotope infusions. E.C.L. was supported by the Van Andel Institute MeNu Program and the NIH (R00CA255928). R.G.J. was supported by the NIH (R01AI165722). This work was also supported by the VAI Core Technologies and Services [Mass Spectrometry (RRID:SCR_024903) and Flow Cytometry (RRID:SCR_022685)].

## Author contributions

E.C.L. and F.C.K. conceived the project. F.C.K., T.J.R., Y.J., A.W., K.H.S., C.J.L., and S.R.D. performed the experiments and analyzed data. T.J.R., J.L., and R.G.J. assisted with generating gene knockout cell lines. A.J. and R.D.S. assisted with mass spectrometry analyses of fatty acids. F.C.K., T.J.R., and E.C.L. wrote the manuscript with input from all authors.

## Competing interests

R.G.J. is a scientific advisor to Servier Pharmaceuticals and is a member of the Scientific Advisory Board of Immunomet Therapeutics. All other authors declare no competing interests.

## Methods

### Cell lines and culture

All human cancer cell lines used for this study were obtained from the American Type Culture Association. AL1376 PDAC cells were isolated from C57BL/6J *LSL-Kras(G12D)*;*Trp53^fl/fl^*;*Pdx1-Cre* mice^4^. B16 cells were obtained from Dr. Russell Jones’ laboratory at the Van Andel Institute. No cell lines used in this study were found in the database of commonly misidentified cell lines that is maintained by the International Cell Line Authentication Committee. Cells were regularly assayed for mycoplasma contamination and passaged for no more than 6 months. All cells were cultured in DMEM (Corning Life Sciences, 10-013-CV) without pyruvate and 10% heat-inactivated dialyzed fetal bovine serum (FBS) (Corning Life Sciences, 35-010-CV) unless otherwise specified. For lipid limitation experiments, FBS was stripped of lipids and dialyzed as previously described to generate lipid-depleted cell culture media^25^.

### Inhibitors

The fatty acid synthase (FASN) inhibitor GSK2194069 (Tocris, 5303) was used at the indicated concentrations.

### Generation of knockout cells by CRISPR-Cas9

sgRNA sequences were chosen from the mouse GeCKO v2 library^26^ and cloned into pSpCas9(BB)-2A-GFP (PX458) (Addgene, #48138). Cells were transfected with these vectors using Lipofectamine 2000 transfection reagent (Invitrogen, 11668027) according to the manufacturer’s protocol. After 48 hours, GFP-positive cells were sorted into single cells with a BD FACSymphony S6 cell sorter and grown up as single-cell clones. Knockouts were confirmed by immunoblotting. sgRNA sequences: *Bdh1 #1,* CGTAGGTCCGACGGGTGTCA*; Bdh1 #2,* AACGCAGGCATCTCAACGTT*; Oxct1* #1, TCTAGGGCACACTTGCCGAG; *Oxct1* #2, ACGAATGATCTCCTCATATG; *Aacs #1*, CGACAGAGTCGCCCTTTACG; *Aacs #2:* TCCGGTCGTATATGGACTTT.

### Immunoblotting

Cells were lysed in radioimmunoprecipitation assay (RIPA) buffer (Thermo Scientific, 89900) supplemented with Halt Protease and Phosphatase Inhibitor Cocktail (ThermoFisher, 78440) for 30 min at 4°C. Cell extracts were pre-cleared by centrifugation at maximum speed for 15 min at 4°C, and protein concentration was measured with the Pierce BCA Protein Assay Kit (Thermo Scientific, 23225). Lysates were resolved on SDS-PAGE and transferred electrophoretically to 0.2 µm nitrocellulose membranes (Bio-Rad, 1620112) at 100 V for 60 min. The blots were blocked in Tris-buffered saline buffer (TBST; 10 mmol/L Tris-HCl, pH 8, 150 mmol/L NaCl, and 0.2% Tween-20) containing 5% (w/v) nonfat dry milk for 30 min, and then incubated with the specific primary antibody diluted in blocking buffer at 4°C overnight. Membranes were washed four times in TBST and incubated with HRP-conjugated secondary antibody for 1 h at room temperature. Membranes were washed three times and developed using SuperSignal West Femto Maximum Sensitivity Substrate (Thermo Scientific, 34096). Antibodies were used as follows: BDH1 (Proteintech, 67448-1-Ig, 1:1000), OXCT1 (Proteintech 12175-1-AP, 1:1000), AACS (Proteintech, 13815-1-AP, 1:2000), Vinculin (Cell Signaling Technology 137015, clone E1E9V, 1:1000), β-actin (Cell Signaling Technology 3700, 1:1000), anti-mouse IgG HRP-linked secondary antibody (Cell Signaling Technology 7076, 1:2000), and anti-rabbit IgG HRP-linked secondary antibody (Cell Signaling Technology 7074, 1:5000).

### Proliferation assays

Cells were seeded at an initial density of 20,000–50,000 cells per well on a 24-well plate in 1 mL of DMEM medium. After incubating for 24 h, cells were washed three times with 1 ml of PBS and changed to the indicated growth conditions. To maintain adequate nutrient levels over time, media was replaced every 24 h over the course of the proliferation assay. Cell confluence was monitored over time using imaging with the Incucyte Live-Cell Analysis System (Sartorius). Doublings per day were calculated by fitting the exponential growth equation to proliferation curves using GraphPad Prism 10.

### Stable isotope labeling experiments and metabolite extraction

Cells were seeded at an initial density of 70,000-150,000 cells per well in a six-well plate in 2 ml of DMEM medium. After incubating for 24 h, cells were washed three times with 2 ml of PBS and then incubated in the indicated media. For glucose isotope labeling experiments, cells were cultured with 10 mM [U-^13^C_6_]-glucose (Cambridge Isotope Laboratories, CLM-1396) for 48 h. For β-OHB isotope labeling experiments, cells were cultured with 5 mM [U-^13^C_4_]-β-OHB (Cambridge Isotope Laboratories, CLM-3853) for 48 h.

For each condition, a parallel plate of cells was scanned with an Incucyte Live-Cell Analysis System (Sartorius) and analyzed for confluence to normalize extraction buffer volumes based on cell number. An empty well was also extracted for a process control. The extraction buffer consisted of chloroform:methanol (containing 25 mg/L of butylated hydroxytoluene (Millipore Sigma, B1378)):0.88% KCl (w/v) at a final ratio of 8:4:3. The final extraction buffer also contained 0.75 µg/ml of norvaline and 0.7 µg/ml of cis-10-heptadecenoic acid as internal standards. For extraction, the medium was aspirated from cells, and cells were rapidly washed in ice-cold saline three times. The saline was aspirated, and methanol:0.88% KCl (w/v) (4:3 v/v) was added. Cells were scraped on ice, and the extract was transferred to 1.5 ml Eppendorf tubes (Dot Scientific, RN1700-GMT) before adding chloroform (Supelco, 1.02444). The resulting extracts were vortexed for 15 min and centrifuged at maximum speed (17000 x g) for 10 min. Polar metabolites (aqueous fraction) were transferred to Eppendorf tubes and dried under nitrogen gas for further analysis. Lipids (organic fraction) were transferred to glass vials (Supelco, 29651-U) and dried under nitrogen gas for further analysis.

### Gas chromatography-mass spectrometry (GC-MS) analysis of fatty acids

Fatty acids were analyzed as pentafluorobenzyl-fatty acid (PFB-FA) derivatives. Fatty acids were saponified from dried lipid pellets by adding 800 µl of 90% methanol/0.3 M KOH, vortexing, and incubating at 80°C for 60 min. Each sample was then neutralized with 80 µl of formic acid (Supelco, FX0440). Fatty acids were extracted twice with 800 µl of hexane and dried under nitrogen gas. To derivatize, fatty acid pellets were incubated with 100 µl of 10% pentafluorobenzyl bromide (Sigma Aldrich, 90257) in acetonitrile and 100 µl of 10% N,N-diisopropylethylamine (Sigma Aldrich, D125806) in acetonitrile at room temperature for 30 min. PFB-FA derivatives were dried under nitrogen gas and resuspended in 50 µl of hexane for GC-MS analysis.

GC-MS was conducted with a TRACE TR-FAME column (ThermoFisher, 260M154P) installed in a Thermo Scientific TRACE 1600 gas chromatograph coupled to a Thermo ISQ 7610 mass spectrometer. Helium was used as the carrier gas at a constant flow of 1.8 ml/min. One microliter of sample was injected at 250°C at a 4:1 split (for total lipid extracts) or splitless mode (for 3PLE extracts). After injection, the GC oven was held at 100°C for 0.5 min, increased to 200°C at 40°C/min, held at 200°C for 1 min, increased to 250°C at 5°C/min, and held at 250°C for 11 min. The MS system operated under negative chemical ionization mode with methane gas at a flow rate of 1.25 ml/min, and the MS transfer line and ion source were held at 255°C and 200°C, respectively. The detector was used in scanning mode with an ion range of 150-500 *m/z*. Total ion counts were determined by integrating appropriate ion fragments for each PFB-FA^27^ using Skyline software^28^. Metabolite data were normalized to the internal standard and background-corrected using a process blank sample. Mass isotopologue distributions were corrected for natural abundance using IsoCorrectoR^29^. Cytosolic acetyl-CoA labeling was calculated from the palmitate mass isotopomer distribution using isotopomer spectral analysis (ISA) from FAMetA^11^.

### Gas chromatography-mass spectrometry (GC-MS) analysis of polar metabolites

Polar metabolites were analyzed as MOX-TBDMS derivatives. Dried and frozen metabolite extracts were derivatized with 16 µl of MOX reagent (Thermo Fisher TS-45950) for 60 min at 37°C, followed by derivatization with 20 µl of N-tert-butyldimethylsilyl-N-methyltrifluoroacetamide with 1% tert-butyldimethylchlorosilane (Millipore Sigma 375934) for 30 min at 60°C. Derivatized samples were analyzed by GC-MS, using a DB-5MS column (Agilent Technologies 122-3832) installed in an Agilent 7890B gas chromatograph coupled to an Agilent 5977B mass spectrometer. Helium was used as the carrier gas at a constant flow rate of 1.2 mL/min. One microliter of sample was injected in split mode (1:4) at 320°C. After injection, the GC oven was held at 95°C for 1 min, increased to 118°C at 40°C/min, held at 118°C for 2 min, increased to 250°C at 12°C/min, ramped to 320°C at 40°C/min, and held at 320°C for 7 min. The MS system operated under electron impact ionization at 70 eV, and the MS source and quadrupole were held at 230°C and 150°C, respectively. The detector was used in scanning mode with an ion range of 50-800 m/z. Total ion counts were determined by integrating appropriate ion fragments for each metabolite using Skyline software^28^. Metabolite data were normalized to the internal standard, and mass isotopologue distributions were corrected for natural abundance using IsoCorrectoR^29^.

### Animal studies

All experiments conducted in this study were approved by the VAI IACUC. For subcutaneous tumor growth, a maximum tumor burden of 2 cm^3^ was permitted per IACUC protocol, and these limits were not exceeded. Male C57BL/6J mice between 3-4 months old were used in this study. All animals were housed at ambient temperature and humidity (18-23°C, 40-60% humidity) with a 12 h light and 12 h dark cycle and co-housed with littermates with ad libitum access to water, unless otherwise stated. All experimental groups were age-matched, numbered, and assigned based on treatment, and experiments were conducted in a blinded manner. Data was collected from distinct animals, where n represents biologically independent samples. Statistical methods were not performed to pre-determine sample size.

For subcutaneous tumors, C57BL/6J mice (The Jackson Laboratory 000664 or internal mouse colony) were subcutaneously injected with 7 × 10^5^ mouse B16 cells into both flanks in 100 µl of phosphate-buffered saline (PBS) per injection. All mice were administered a modified AIN-93G control diet (Envigo TD.97184) during tumor formation, and 7-9 days after cell injection when palpable tumors had formed, animals were randomly placed into different diet groups. Mice were weighed before the start of diet administration to ensure different cohorts had similar starting body weights, and body weights were also measured over the course of each experiment. Tumor volume was determined using (π/6)(*W* ^2^)(*L*), where *W* represents width and *L* represents length as measured by calipers. At the end of each experiment, animals were euthanized, blood was collected by orbital bleed, and tumors or tissues were rapidly harvested, weighed, and freeze-clamped in liquid nitrogen.

### *In vivo* [U-^13^C]-β-OHB infusions

*In vivo* infusions of [U-^13^C]-β-OHB were conducted using previously described protocols^23^. Briefly, tumor-bearing mice were fasted for 16 hours before being infused with [U-^13^C]-β-OHB (Cambridge Isotope Laboratories, CLM-3853) through a tail vein catheter. Mice received an initial bolus of 478 mg/kg for 1 minute followed by infusion of 0.05 μL/min/g using a 1.5 M stock concentration for 6.5 hours. Subsequently, mice were cervically dislocated, and tumors were freeze-clamped in liquid nitrogen. Blood was harvested via orbital bleed, mixed with EDTA, centrifuged at 3000 RCF for 10 min at 4°C, and plasma was snap frozen in liquid nitrogen. Frozen tumors were crushed into a fine power using a chilled mortar and pestle and stored at - 80°C. 5-20 mg of powdered tumor were measured for metabolite extraction.

### Animal diets

A modified AIN-93G diet (Envigo TD.97184) was used as the control diet. The 40% caloric restriction (CR) diet (Envigo TD.210722) was formulated by modifying the control diet, as previously described^4^. For CR studies, mice were individually housed. Prior to diet administration, the average daily consumption (by weight) of the control diet was determined. Upon experimental diet initiation, control mice were fed daily with the determined average daily food consumption weight, and calorie restricted mice were fed daily with a food weight corresponding to 40% of the control caloric consumption.

### Statistics and reproducibility

Sample sizes, reproducibility, and statistical tests used for each figure are denoted in the figure legends. All graphs were generated using GraphPad Prism 10.

### Data availability

All data generated and analyzed during this study are included in this published article. Correspondence and requests for materials should be addressed to Evan C. Lien.

## Extended Data Figure Legends

**Extended Data Figure 1.**
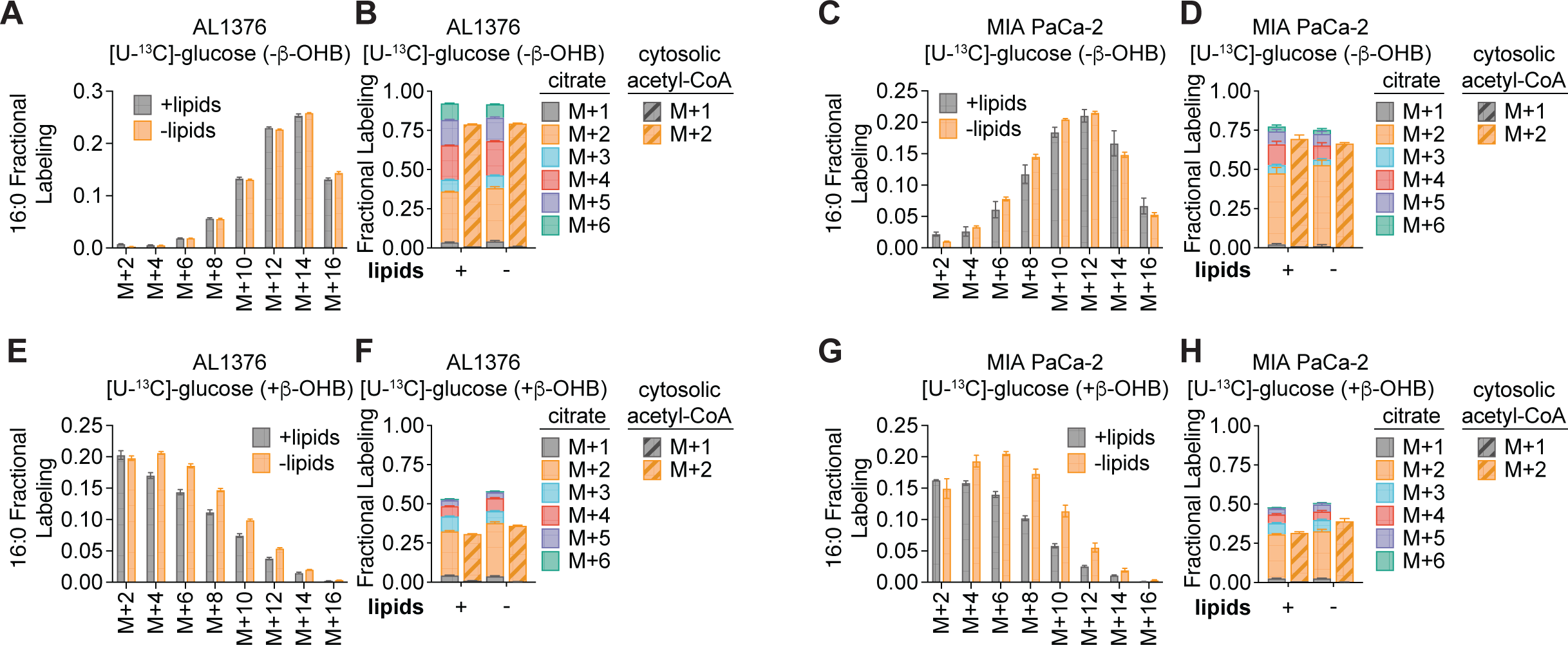
β-OHB is preferentially used over glucose for fatty acid synthesis, Related to Figure 2. **A-D**, 16:0 MID (**A**, **C**) and citrate (solid bars) and cytosolic acetyl-CoA (dashed bars) MID (**B, D**) for AL1376 (**A**, **B**) and MIA PaCa-2 (**C**, **D**) cells labeled with [U-^13^C]-glucose for 48 h in lipid-replete versus lipid-depleted culture media without unlabeled β-OHB. **E-H**, 16:0 MID (**E**, **G**) and citrate (solid bars) and cytosolic acetyl-CoA (dashed bars) MID (**F, H**) for AL1376 (**E**, **F**) and MIA PaCa-2 (**G**, **H**) cells labeled with [U-^13^C]-glucose for 48 h in lipid-replete versus lipid-depleted culture media with 5 mM unlabeled β-OHB. Data are presented as mean ± s.e.m; *n* = 3 biologically independent replicates.

**Extended Data Figure 2.**
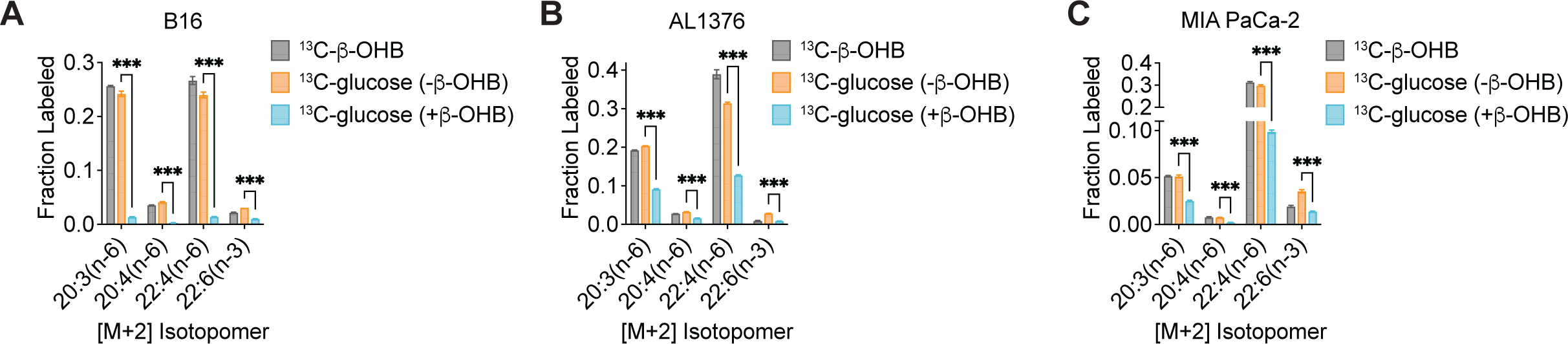
β-OHB is preferentially used over glucose for PUFA elongation, Related to Figure 2. **A-C**, [M+2] fractional labeling of 20:3(n-6), 20:4(n-6), 22:4(n-6), and 22:6(n-3) in B16 (**A**), AL1376 (**B**), and MIA PaCa-2 (**C**) cells labeled with [U-^13^C]-β-OHB, [U-^13^C]-glucose without unlabeled β-OHB, or [U-^13^C]-glucose with 5 mM unlabeled β-OHB for 48 h in lipid-replete media. Data are presented as mean ± s.e.m; *n* = 3 biologically independent replicates. Comparisons were made using a two-tailed Student’s t test (**A-C**). \*\*\**P<0.001*.

**Extended Data Figure 3.**
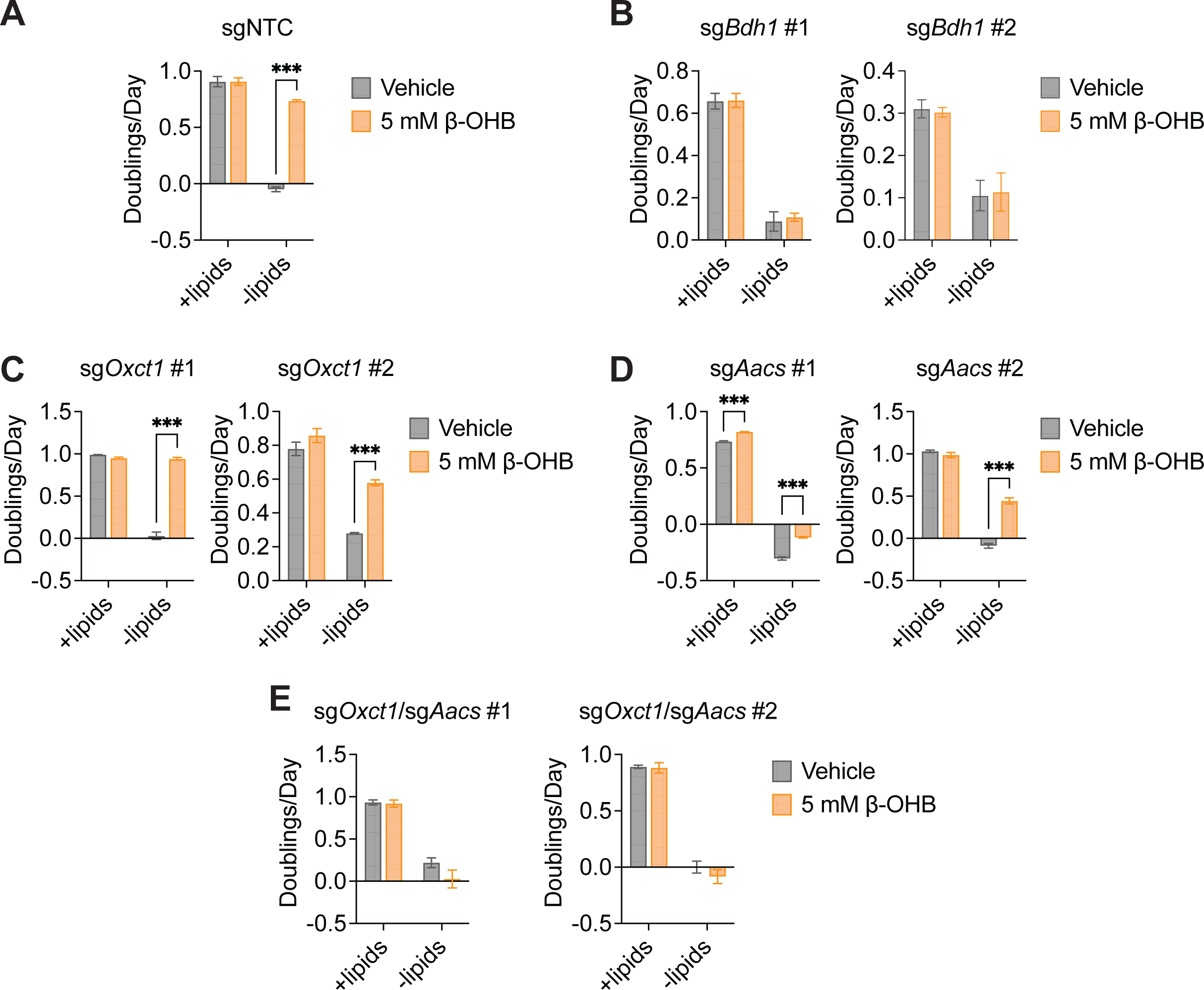
β-OHB can contribute to fatty acid synthesis through a citrate-independent route that requires AACS, Related to Figure 3. **A-E**, Proliferation rates of the indicated B16 sgNTC (**A**), sg*Bdh1* (**B**), sg*Oxct1* (**C**), sg*Aacs* (**D**), and sg*Oxct1*/*Aacs* (**E**) cells grown in lipid-replete versus lipid-depleted culture media, with or without 5 mM β-OHB. Data are presented as mean ± s.e.m; *n* = 3 biologically independent replicates. Comparisons were made using a two-way ANOVA. \*\*\**P<0.001*.

**Extended Data Figure 4.**
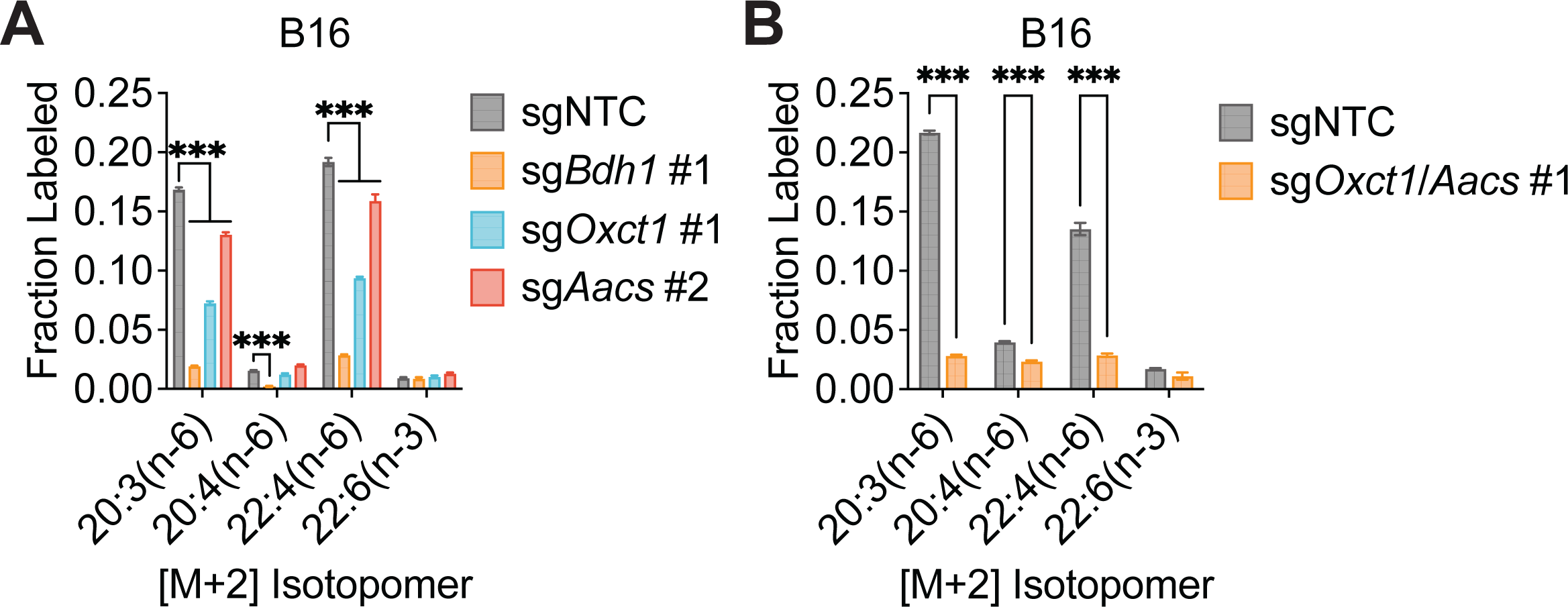
β-OHB contributes to PUFA elongation through both OXCT1 and AACS, Related to Figure 3. **A**, [M+2] fractional labeling of 20:3(n-6), 20:4(n-6), 22:4(n-6), and 22:6(n-3) in B16 sgNTC, sg*Bdh1* #1, sg*Oxct1* #1, and sg*Aacs* #2 cells labeled with [U-^13^C]-β-OHB for 48 h in lipid-replete media. **B**, [M+2] fractional labeling of 20:3(n-6), 20:4(n-6), 22:4(n-6), and 22:6(n-3) in B16 sgNTC and sg*Oxct1*/*Aacs* #1 cells labeled with [U-^13^C]-β-OHB for 48 h in lipid-replete media. Data are presented as mean ± s.e.m; *n* = 3 biologically independent replicates. Comparisons were made using a two-way ANOVA. \*\*\**P<0.001*.

**Extended Data Figure 5.**
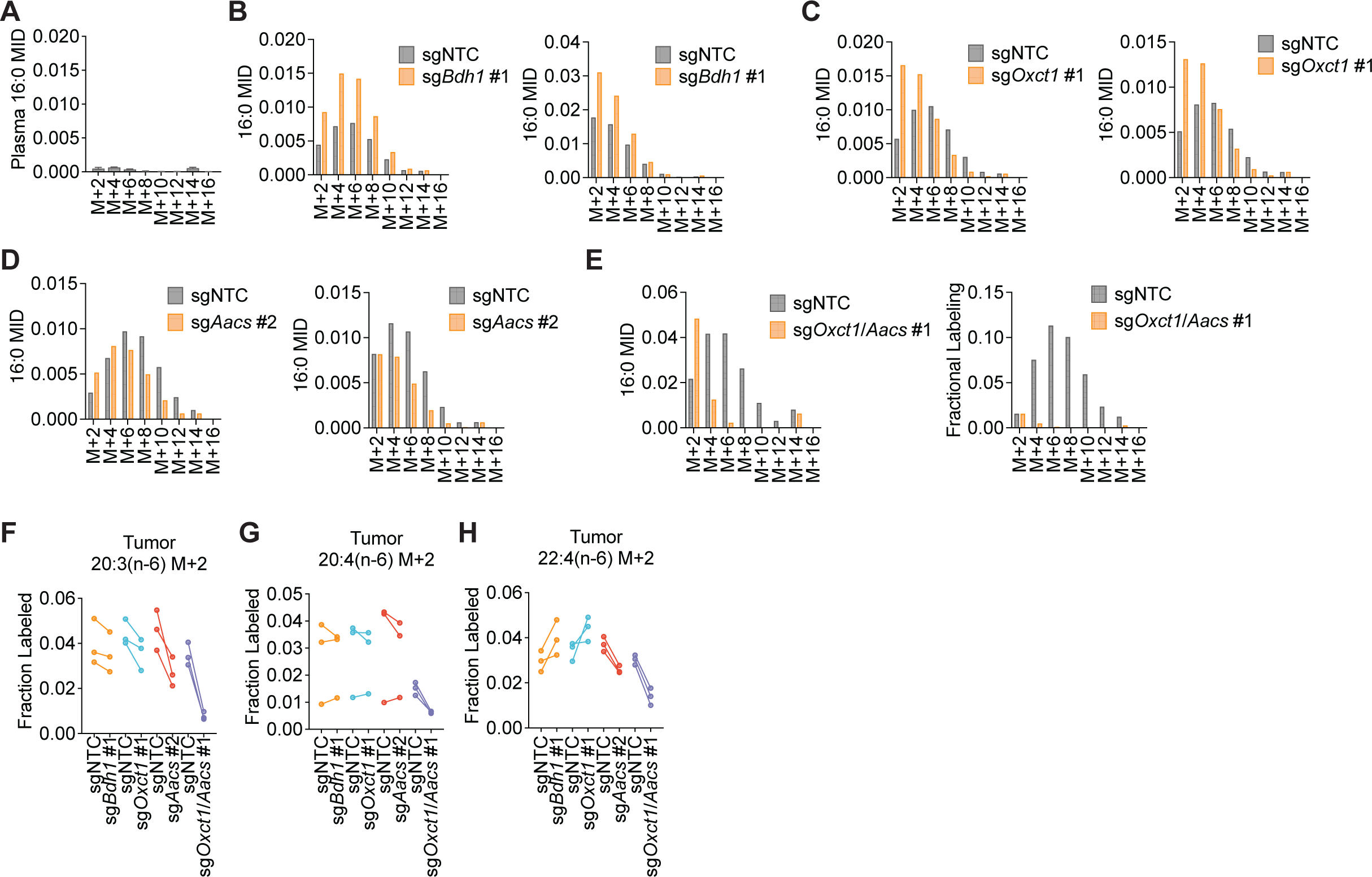
β-OHB contributes to fatty acid synthesis through both OXCT1 and AACS in tumors *in vivo*, Related to Figure 4. C57BL/6J mice bearing a B16 sgNTC tumor on one flank and a knockout tumor on the other flank were infused with [U-^13^C]-β-OHB for 6.5 h. **A**, Plasma 16:0 MID in all mice. **B-E**, 16:0 MIDs from each mouse bearing an sgNTC tumor versus a sg*Bdh1* #1 tumor (**B**), sg*Oxct1* #1 tumor (**C**), sg*Aacs* #2 tumor (**D**), and sg*Oxct1*/*Aacs* #1 tumor (**E**). **F-H**, [M+2] fractional labeling of 20:3(n-6) (**F**), 20:4(n-6) (**G**), and 22:4(n-6) (**H**) in the indicated sgNTC versus knockout tumors. Data are paired between sgNTC and knockout tumors from the same mouse. n = 3 biological replicates for each knockout tumor. Data are presented as mean ± s.e.m.

